# Single cell RNA sequencing reveals limited effects of Lrrk2 genotype in a mouse model of acute CNS inflammation

**DOI:** 10.64898/2025.12.18.695199

**Authors:** Alice Kaganovich, Liam Horan-Portelance, Natalie Landeck, Dominic J Acri, Jinhui Ding, Rebekah Langston, Mark R Cookson

## Abstract

**BACKGROUND:** Previous data implicates neuroinflammation in the pathogenesis of Parkinson’s disease (PD). Of the genes associated with PD, Leucine rich repeat kinase 2 (LRRK2) has previously been proposed to play a role in neuroinflammation. However, the extent to which LRRK2 is involved in endogenous inflammatory signaling is unclear.

**OBJECTIVE:** To examine whether endogenous mutations in LRRK2 affect responses to neuroinflammation *in vivo*.

**METHODS:** We injected cohorts of mice with knock-in mutations in *Lrrk2*, homologous to those causing human PD, with a single intrastriatal injection of lipopolysaccharide (LPS) or control. We used single cell RNA-Sequencing to examine cell type specific responses to treatment and genotype and validated key results with orthogonal approaches.

**RESULTS:** We found that our chosen paradigm of acute LPS exposure evokes robust transcriptional changes consistent with a multicellular neuroinflammatory response. We also found evidence of peripheral immune cell recruitment into the brain and interaction with brain-resident microglia. However, the transcriptional effects of Lrrk2 mutations were limited to small numbers of genes, including down regulation of gene ontogeny terms related to lysosomes, in microglia.

**CONCLUSIONS:** Our data clearly demonstrate that many cells in the brain respond to a single inflammatory insult with strong transcriptional responses and that, even in a model focused on CNS injection, there is interaction between peripheral and central immune cells. In contrast, the quantitative effects of Lrrk2 mutations are modest at the transcriptional level, demonstrating that additional studies are needed to clarify whether Lrrk2 mutations affect neuroinflammation in an endogenous context.

## Introduction

Age-related neurodegenerative diseases including Parkinson’s disease (PD) are characterized in part by selective neuronal cell loss and variable protein deposition pathology [1]. Additionally, neuroinflammation is a prominent neuropathological event in affected brain areas in PD [2]. Inflammatory reactions in PD encompass both a transition of microglia [3] and astrocytes [4] to their various reactive states as well as infiltration of external cells from the peripheral immune system, including neutrophils and macrophages [5]. It is likely that this extensive increase in neuroinflammation is a result of cross-talk between multiple cell types and may change as PD pathology evolves over time [2].

Whether neuroinflammation is a primary or secondary factor in neurodegeneration remains unclear [6]. On the one hand, because glia and other immune cells are primed to react to cellular damage, neuroinflammation may occur as a sequelae of neuronal dysfunction including protein aggregation and release from damaged cells [7]. On the other hand, depletion of microglia or macrophages has been shown to be neuroprotective in some PD models, suggesting that these cells make an active contribution to neuronal damage [8–10]. However, microglial depletion may also be detrimental in some contexts [11,12].

One argument in favor of neuroinflammation being a causative factor in neurodegeneration is that several genes associated with neurological disorders are expressed in non-neuronal cells. For example, autosomal dominant mutations in *Leucine-rich repeat kinase 2 (LRRK2)* cause familial PD [13–15] and the gene product is found in a subset of neurons [16] but also in peripheral immune cells [17,18] and microglia [19]. Furthermore, we have recently demonstrated that LRRK2 levels may contribute to sporadic PD risk by virtue of a microglial-specific expression quantitative trait locus (eQTL) [20], consistent with prior data in peripheral monocytes [21]. Cells deficient in LRRK2 show diminished inflammatory reactions across a wide range of stimuli, notably involving toll-like receptor (TLR) signaling [22–24]. At a cellular level, LRRK2 has important roles in control of organelle membranes via RAB proteins [25] and has been specifically linked to lysosomal membrane repair [26–29]. This pathway is intimately linked to inflammation as, for example, activated microglia increase uptake of debris and lysosomal turnover [30–33]. These data collectively suggest that LRRK2 may play a causal role in inflammatory signaling.

Prior studies also indicate a role for Lrrk2 in the progression of inflammation *in vivo* models. For example, several studies utilizing *Lrrk2* knockout mice or kinase inhibitors have shown a blunted immune response to inflammatory models, including α-synuclein spread [34,35] or LPS exposure [36]. Additionally, mice carrying *LRRK2* mutations expressed from a bacterial artificial chromosome transgene have shown alterations in peripheral and central immune signaling, mediated by peripheral lymphocytes [37,38]. These data collectively suggest that LRRK2 plays a modulating role in inflammation with the prediction that autosomal dominant mutations associated with PD would be pro-inflammatory. However, these data have not resolved which CNS cell types, if any, contribute to the effects of LRRK2 in the brain. Specifically, the extent to which resident microglia, peripheral immune cells or even astrocytes or neurons are affected by LRRK2 mutations is uncertain.

To attempt to address the problem of which cells in the brain mediate effects of LRRK2, we used single cell RNA-Sequencing to examine transcriptional responses to an acute neuroinflammatory stimulus. To provide an endogenous context, we chose two mouse models where the equivalent mutation to human LRRK2 was knocked into the mouse genome, R1441C [39] and G2019S [40]. We also chose to use a single intrastriatal injection of lipopolysaccharide (LPS) and evaluated whether this treatment paradigm induced expected changes in gene expression in local immune cells and whether there was evidence of peripheral cell infiltration in a model where inflammation was initiated within the brain. While we were able to conclusively demonstrate and validate that there was a strong response of multiple cell types to LPS exposure, the effects of Lrrk2 genotype were surprisingly subtle, limited to a few gene expression changes related to lysosomal function. Overall, our results suggest that Lrrk2 is not a major driver of inflammation-induced transcriptional responses.

## Materials and Methods

### Animals

Mice were housed in standardized conditions with 2 to 5 animals per cage on a 12-hour light/dark cycle with food and water provided *ad libitum*. C57BL/6J (WT) was obtained from Jackson Laboratory (Jax) (strain #000664). Lrrk2 p.G2019S knock-in mice (B6.Cg-LRRK2^tm1.1Hlme^/J) [40] and were obtained from Jax (strain #030961). Lrrk2 R1441C knock-in mice (B6.Cg-LRRK2^tm1.1Shn^/J) [39] and were also obtained from Jax (strain #009346). Each strain was backcrossed to the WT line for a minimum of 2 generations. All protocols used here were approved by the Institutional Animal Care and Use Committee (ACUC) of National Institute on Aging, NIH (protocol ID: 463-LNG-2024) and strictly followed.

### Stereotaxic Surgery

12- to 14-month-old male and female mice (n = 3 per group) were anesthetized using 5% isoflurane and kept under plane of anesthesia using 1–2% isoflurane. Mice were placed into a stereotaxic frame (Kopf Instruments) and eyes were covered with eye ointment (CVS). The tooth bar was adjusted to −5.0 mm. The top of the head was shaved and sterilized using 70% ethanol and betadine. An incision was made above the midline and the skull was exposed using cotton tips. A hole was drilled into the skull at anteroposterior +0.2 mm, mediolateral ± 2.0 mm from bregma (bilateral injection), and the last layer of the skull was removed using a forceps to not damage the dura. A pulled glass capillary attached to a 5 μl Hamilton syringe was used to inject 1 μl of either PBS or 5mg/ml LPS dissolved in PBS (E.coli, serotype O111:B4, Sigma). The capillary was lowered to dorsoventral −3.2 mm from bregma into the dorsal striatum. The solution was delivered at a rate of 0.1 μl per 10 sec. After injection, the capillary was held in place for 2 min, retracted 0.1 μm and, after another 1 minute, was slowly withdrawn from the brain. The head wound was closed using surgical staples. Ketoprofen solution at 5 mg/kg was administered subcutaneously as analgesic treatment for the following 48 hours.

### 10x Genomics single-cell RNA sequencing

Forty eight hours post-injection, animals were euthanized using CO2. Brains were quickly removed, washed in ice-cold PBS, and dissected on ice using a mouse brain matrix. Striatal and ventral midbrain tissue samples of each animal were combined into one sample and dissociated together using the Adult Brain Dissociation kit (Miltenyi Biotech, Cat #130-107-677) followed by Myelin removal using the Myelin Removal Beads (Miltenyi Biotech, cat #130-096-731) according to the manufacturer’s recommendation.

Single cell sequencing libraries were then prepared from 5000 cells using the Chromium Next GEM Single Cell 3’ Reagent Kits v3.1 (10x Genomics #PN-1000121). Libraries were sequenced to a paired-end depth of 100,000 reads per cell on a NovaSeq 6000 using the Flow Cell S4 (Illumina) at the NIH Intramural Sequencing Center.

### scRNA-seq data analysis

Illumina sequencing reads were first preprocessed using the 10x Genomics CellRanger pipeline, which was used to align reads to the mouse mm10 reference genome and count transcripts. After preprocessing, samples were independently quality controlled by removing genes with low expression (number of cells expressing a gene > 3) and low-quality cells (number of unique genes expressed > 500, percent mitochondrial genes < 20). Of note, one animal in the Lrrk2 G2019S + LPS treatment group was excluded from the analysis due to poor condition of the mouse and the absence of a classical inflammatory response in the associated scRNA-seq data.

Samples that passed quality control were then integrated using the Seurat package (v5) in R [41]. Datasets were first independently normalized using the SCTransform function while regressing out percent mitochondrial genes. PCA was computed on the first 30 dimensions, and samples were integrated using the *IntegrateLayers* function, with “method” set to “RPCAIntegration”. We then ran the standard Seurat workflow on the integrated object (*RunUMAP*, *FindNeighbors*, *FindClusters*) and identified 16 major clusters of cells at a clustering resolution of 0.2. The top marker genes from each cluster were computed using the *FindAllMarkers* function in Seurat, and cell types were manually identified and assigned using canonical cell type markers.

For subcluster analysis, cell types of interest were subset out of the main object, and the standard Seurat workflow was run on the subset object as above. Statistical inferences for cell proportions on a per animal basis were performed using two-way ANOVA for genotype and treatment, with Tukey’s multiple comparison’s test for individual comparisons if the parent ANOVA was significant, performed in GraphPad Prism.

### Pseudobulk differential expression analysis

For examining broad differences in gene expression within cell types, we opted for a pseudobulk approach, for which we used DESeq2 [42]. When creating our design matrix, we assigned genotype (WT, RC, or GS) and treatment (PBS or LPS) as variables, with an interaction term (∼genotype + treatment + genotype:treatment). We then performed differential expression, comparing PBS vs. LPS-treated mice, as well as comparing LPS-treated mice of different genotypes using the interaction term. For identifying differentially-expressed genes, we considered genes with a log_2_FC > 0.25 or < −0.25 and a FDR p-value < 0.1 to be significant, given our relatively small n of 2 or 3 animals per group.

### Cell-cell communication analysis

For cell-cell communication analysis, we utilized CellChat following the standard workflow. Briefly, different genotype x treatment conditions (e.g., WT_PBS, WT_LPS, etc.) were normalized and cell-cell interaction networks were predicted separately.

Following cell-cell interaction network prediction, the outgoing and incoming signals predicted within microglia from LPS-treated brains were calculated.

### RNAscope In situ hybridization

A separate cohort of WT animals underwent LPS or PBS striatal injections to confirm scRNAseq data using RNAscope in situ hybridization (ISH) by HiPlex or Multiplex RNAscope. 48 hr after injection, mice were deeply anesthetized using an overdose of sodium pentobarbital (30 mg/kg). When reflexes were absent, the animal was transcardially perfused using ice cold PBS for 2 min. The brain was quickly taken out and cut into two hemispheres. One hemisphere was flash frozen for ISH. The other hemisphere was dissected for striatal and midbrain areas, flash frozen and stored at −80 C for future use.

The frozen whole hemisphere was fully embedded in O.C.T. compound (Tissue-Tek). Each block was cryosectioned into 16um-thick coronal slices using a Leica CM3050 Cryostat, mounted onto positively charged slides (Fisher Brand, Cat #12-550-15) and stored at −80C.

ISH was performed using either HiPlex12 Reagent Kit (Cat #324410, Advanced Cell Diagnostics (ACD), Newark, CA, USA) or Multiplex Fluorescent Detection kit (Cat #323110, ACD) according to the manufacturer’s protocols. Briefly, sections were fixed with 4% paraformaldehyde (60 min for HiPlex and 15 min for Multiplex RNAscope), treated with either Protease III or IV for 25 min and subjected to 2.5 min of boiling in the target retrieval reagent (Cat #322000, ACD) followed by the HiPlex v2 or the Multiplex v2 RNA scope protocols. For detection of the Multiplex RNA scope probes, the compatible Opal fluorophores were purchased separately from Akoya Biosciences (Cat #FP1487001KT for Opal 520, Cat #FP1488001KT for Opal 570 and Cat #FP1496001KT for Opal 650). Each probe was tested in 2-4 sections from at least n=2 animals.

The following gene-specific RNAscope HiPlex probes have been used as identified from scRNAseq analysis:

Cat #313971-T4 RNAscope HiPlex Probe - Mm-Lcn2-T4,

Cat #446841-T7 RNAscope HiPlex Probe - Mm-Saa3-T7

Cat #511561-T3 RNAscope HiPlex Probe - Mm-Fth1-T3

Cat #319141-T2 RNAscope HiPlex Probe - Mm-Aif1-T2

Cat #529541-T8 RNAscope HiPlex Probe - Mm-Bin1-T8

Cat #824201-T9 RNAscope HiPlex Probe - Mm-Msr1-T9

For quality control, tissues were incubated with positive and negative species-specific control probes to evaluate tissue and RNA quality.

### Microscopy for RNA scope ISH

For Multiplex RNAscope and after each round of the HiPlex RNAscope, images were obtained using a Zeiss LSM780 laser scanning microscopy. A 20x objective has been used to acquire the whole striatum (tiling combined with z-stacks) and 40x or 63x oil immersion objectives have been used to obtain a representative high magnification image. Imaging tiles settings of 20x magnification were stitched using Zeiss ZEN software and maximum intensity projections (MIP) were generated. After each round of imaging, cover slips were removed, fluorophores were cleaved with the Cleaving Solution (Cat #324130, ACD) according to manufacturer’s protocol and sections were incubated with new fluorophores to detect the next set of probes in the same cells on the same tissue slices.

### Image analysis and quantification of RNAscope puncta

Images of the whole striatum were split between channels and batch exported using Zeiss Zen software (blue edition) and registered using RNAscope HiPlex Image Registration Software v2.0 (Advanced Cell Diagnostics (ACD), Newark, CA, USA) The resulting ome.tif files were uploaded and analyzed in QuPath (Open Source Software for Bioimage Analysis).

Consistent striatal area were annotated for each set of PBS and LPS treated samples and DAPI-stained nuclei were detected. Positive Percent Area for each specific probe was calculated based on RNA puncta signal according to each gene’s threshold cutoff and exported as “Annotation measurement” for each set of the treated sections.

### Histology

For immunohistochemistry, a fresh set of animals was treated with LPS or PBS and euthanized 48 hr after surgery. Animals were deeply anesthetized using an overdose of sodium pentobarbital (30 mg/kg). Mice were perfused via the ascending aorta first with 10 ml of 0.9% NaCl (2 min) followed by 50 ml of ice-cold 4% paraformaldehyde (PFA in 0.1 mM phosphate buffer, pH 7.4) for 5 min. Brains were removed and post-fixed in 4% PFA for 24 h and then transferred into 30% sucrose for cryoprotection. The brains were then cut into 30 µm thick coronal sections (horizontal freezing microtome, Leica) and stored in an antifreeze solution (0.5 mM phosphate buffer, 30% glycerol, 30% ethylene glycol) at –20°C until further processed.

For astrocyte evaluation, striatal sections were rinsed with PBS and incubated for 30 minutes in blocking buffer (10% Normal Donkey Serum (NDS), 1% BSA, 0.3% Triton in PBS). Afterwards, primary antibody rabbit anti-Gfap (ab7260, abcam) was applied at 1:500 and incubated overnight at room temperature in 1% NDS, 1% BSA, 0.3% Triton in PBS. Next day, sections were rinsed 3x for 10 minutes each with PBS and incubated with Alexa Fluorophore 568-conjugated secondary antibodies for 1 hour at room temperature. After 3 washes with PBS, sections were mounted on glass slides, coverslipped using Prolong Gold Antifade mounting media (Invitrogen), and imaged using a Zeiss LSM 880 confocal microscope equipped with Plan-Apochromat 63X/1.4 numerical aperture oil-objective (Carl Zeiss AG).

## Results

### Single cell analysis of LPS exposure in the central nervous system

For the current experiments, we injected LPS or PBS as vehicle into the striatum of wild type C57bl/6J, Lrrk2 p.R1441C [39] or Lrrk2 p.G2109S [40] mice, leading to activation of cells throughout the mouse brain. Using single cell RNA-seq (scRNA-seq), we then examined transcriptional responses to neuroinflammation in the striatum and ventral midbrain, 2 days after injection [Fig. 1A]. We projected 76,305 recovered cells using UMAP [Fig. 1B; Table S1] and classified cell types based on canonical gene markers [Fig. 1C; Table S2]. We were able to reliably identify both neuronal and non-neuronal cells, including endothelial cells ([Endo] *Cldn5*), peripheral myeloid cells (*Ncf4, Cd44*), red blood cells ([RBC] *Hbb*), astrocytes ([Astro] *Aqp4*), microglia ([Micro] *Tmem119*), immature neurons (*Dcx*, *Sox11*), pericytes and vascular smooth muscle cells ([Peri_VSMC] *Vtn*), oligodendrocytes and oligodendrocyte precursors ([Oligo_OPC] *Plp1*, *Pdgfra*), lymphocytes (*Cd3e*), neurons (*Rbfox3*, *Snap25*), including spiny projection neurons ([SPN] *Gad1, Ppp1r1b*), choroid plexus cells (*Ttr*), ependymal cells (*Foxj1*), vascular leptomeningeal cells ([VLMC] *Dcn*), fibroblasts (*Prg4*), and tancycytes (*Col23a1*).

**Figure 1.**
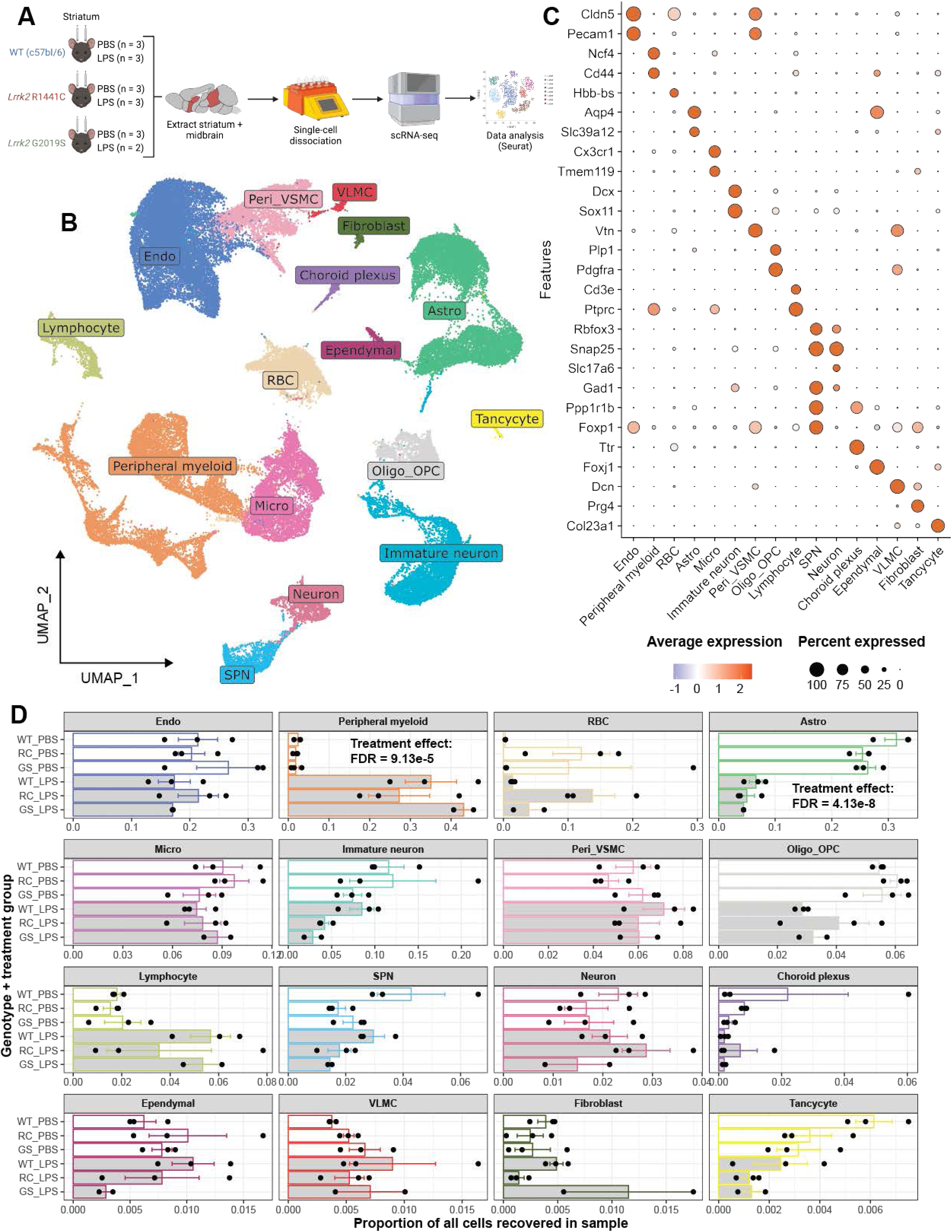
**Single-cell analysis of an acute model of central nervous system inflammation.** [A] Experimental design. Wild type (C57Bl/6J; WT) or two knockin mutations in mouse Lrrk2 (p.R1441C and p.G2019S) equivalent to those associated with human Parkinson’s disease were injected with PBS as vehicle or a single dose of LPS in the striatum. Dissected striatum and ventral midbrain areas were combined, dissociated to single cells and analyzed using scRNA-seq. [B,C] Cell types visualized using Universal Manifold Approximation and Projection (UMAP) to show cell clusters for all animals and treatments, labeled by inferred cell type [B], which is based on known cell types as indicated in the dotplot [C] sized by percentage of cells with detectable expression and colored by average expression levels in each cell cluster. See Table S1-2 for cell taxonomy and marker statistics. [D] Proportions of major cell types (colored as in A) across combined treatments (open bars are PBS and gray bars are LPS) and genotypes by sample. Each dot represents a single animal. Two-way ANOVA with Tukey’s multiple comparisons (Table S3; *p < 0.05, **p < 0.01, ***p < 0.001, ****p < 0.0001 comparing LPS and PBS treated animals for the same genotype; no genotype differences were significant).

In general, all cell types were found in both treated and untreated animals as well as all genotypes with three exceptions. First, peripheral myeloid cells were predominantly recovered from LPS- over PBS- treated animals [Fig. 1D; Table S3], suggesting that these cells were recruited from the periphery towards the brain (two-way ANOVA; F_1,11_ treatment = 82.27, B-H FDR = 9.12E-5). Second, we noted poorer recovery of astrocytes in LPS- compared to PBS-treated samples (two-way ANOVA; F_1,11_ treatment = 366.0, B-H FDR = 4.13E-8). Given that astrogliosis is a prominent reaction to neuroinflammation [4], we considered whether this observation may result from artificial (rather than biological) differences in recovery of whole cells during isolation prior to sequencing. Using immunostaining, we noted a clear upregulation of glial fibrillary acidic protein (Gfap), a marker of astrocyte activation [4], in the striatum of LPS treated animals [Supplementary Fig. 1], indicating that the loss of astrocytes thus likely represents paucity of recovery of activated cells after treatment. Finally, the cluster containing oligodendrocytes and OPCs (Oligo_OPC) displayed a significant treatment effect (two-way ANOVA; F_1,11_ treatment = 25.12, B-H FDR = 1.82E-2). However, because we performed myelin removal prior to library preparation, we did not consider these measurements to be reliable and did not further analyze this cluster. Outside of these exceptions, cells were recovered equally across treatment groups, and there were no significant genotype or genotype:treatment interaction effects. Hence, we proceeded to perform gene expression analysis within subtypes of these major cell types.

### A single intrastriatal injection of LPS results in peripheral immune cell infiltration into the brain

To analyze the cell type-specific response to intrastriatal LPS exposure, we first sought to understand from which cells the inflammatory cascade might originate. Accordingly, we analyzed the expression patterns of the two primary receptors for LPS, *Tlr4* and *Cd14*, in the PBS-treated samples only, as to not confound the influence of LPS on the expression of these genes. In accordance with prior literature [43], these receptors were predominantly expressed in immune cells, microglia and macrophages [Supplementary Fig. 2]. Therefore, we first analyzed gene expression in immune cells in this model.

Subclustering of brain-resident and peripheral myeloid cells followed by UMAP projections of microglia (MG), neutrophils (NP), monocytes (MO), and macrophages (MØ) demonstrated strong changes in proportions of each cell type between PBS and LPS treated animals [Fig. 2A; Table S4]. We further divided each cell type into transcriptionally distinct cell states. We were able to separate microglia into two populations. Homeostatic microglia (Micro_homeo) with higher expression of resting markers such as *P2ry12* and *Tmem119* were recovered mainly in PBS-treated animals [Fig. 2A,B; Table S5] (two-way ANOVA; F_1,13_ treatment = 41.63, B-H FDR = 2.84E-4). In contrast, a subset of inflammatory reactive microglia (Micro_inflamm) were enriched with LPS exposure [Fig 2A, B] (two-way ANOVA; F_1,13_ treatment = 41.63, B-H FDR = 2.84E-4). This group of cells were characterized by high expression of cytokines and chemokines including *Ccl5*, which has been previously shown to be expressed by activated microglia in PD and regulate interactions with T cells [44]. This population additionally strongly expressed complement genes (e.g., *C1qa/b/c*) and ribosomal genes (e.g., *Rpl/Rps* genes) [Fig 2B]. While these results indicate that there are substantial changes in gene expression in microglia in response to LPS *in vivo*, we did not see any significant differences in cell proportion between *Lrrk2* genotypes at baseline or in an interaction model with treatment [Fig. 2C; Table S6].

**Figure 2.**
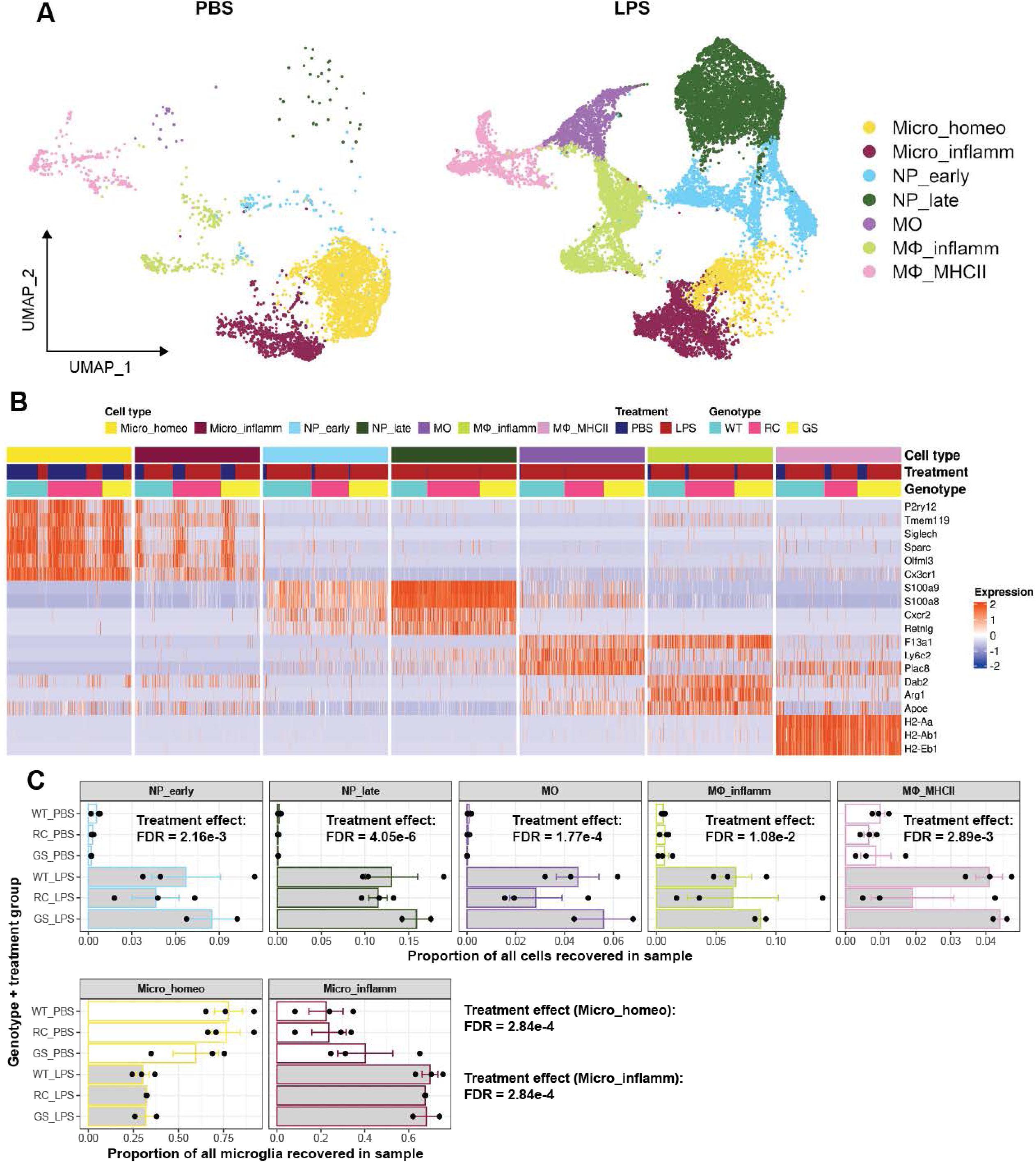
**Changes in transcriptional state of immune cells after intrastriatal LPS injection.** [A] Universal manifold approximate projections of immune cell subsets in PBS (left) and LPS (right) treated mice. MG, microglia; NP, neutrophil; MO, monocytes; Mɸ, macrophage. See Table S1 for immune cell taxonomy. [B] Heatmap of selected gene expression markers in immune cell subsets (Cell types colored as in A, followed by treatment and genotype) showing single cell expression of representative marker genes for each cell type and state. See Table S2 for markers. [C] Proportions of major cell types (colored as in A) across combined treatments (open bars are PBS and gray bars are LPS) and genotypes by sample. Each dot represents a single animal. Two-way ANOVA with Tukey’s multiple comparisons (Table S3; *p < 0.05, **p < 0.01, ***p < 0.001, ****p < 0.0001 comparing LPS and PBS treated animals for the same genotype; no genotype differences were significant).

In contrast to brain-resident microglia, the proportion of cells identified as neutrophils, monocytes or macrophages were low in PBS-treated animals and were all significantly higher after LPS exposure [Fig. 2A]. Furthermore, we were able to separate both neutrophils and macrophages into two transcriptionally distinct states. Neutrophils were separated into ‘early’ and ‘late’ responding cells due to the higher expression of *S100a8* and *S100a9* in the latter population, which are known markers of inflammation [45].

Finally, we found two states of macrophages, one characterized as “inflammatory” (MØ_inflamm) by markers such as *Arg1*, *Dab2*, and *Apoe*, and one state with high expression of MHC-class II genes (MØ_MHCII) [Fig. 2B]. These results show that a single intra-striatal LPS exposure results in extensive recruitment of multiple inflammatory cell types to the brain. However, *Lrrk2* genotype again did not influence the proportion of cells that were recovered after LPS exposure [Fig. 2C].

### Validation of common and cell type specific gene expression changes in response to LPS

We next examined shared and cell specific expression changes in response to LPS and validated a set of genes using the orthogonal approach of HiPlex RNAscope. We identified *Lcn2*, which is known to play a role in innate immunity by sequestering iron [46], as being upregulated strongly across many cell types [Supplementary Fig. S3B]. We were able to show general upregulation of Lcn2 in the striatum using RNAScope [Supplementary Fig. S3A].

Similarly, Saa3, which enables TLR4 signaling [47] was generally upregulated in single cell analysis [Supplementary Fig. S3F] and was validated by RNAscope [Supplementary Fig. S3E]. Additionally, we examined *Fth1* expression, which we have previously nominated as a gene responsive to LPS exposure in isolated microglia [48]. The current single cell data confirmed prior work and extended to show that all cells show upregulation of *Fth1* [Supplementary Fig. S3D] that, again, was validated using RNAScope [Supplementary Fig. S3C].

We next attempted to validate changes that were restricted to specific cell types. For example, *Bin1* is expressed in multiple cell types and shows decreased expression in LPS exposed animals [Supplementary Fig. S4B]. In contrast, macrophage scavenger 1 (*Msr1*) was increased in microglia [Supplementary Fig. S4C]. Using *Aif1*, which codes for the protein Iba1 that is widely used as a marker for microglia, as a reference, we were able to show dramatic downregulation of *Bin1* and upregulation of *Msr1* in LPS treated animals [Supplementary Fig. S4A]. These data collectively validate gene regulation changes in response to LPS across cell types *in vivo*.

### LPS exposure induces changes in genes involved in cell-cell communication

In order to better understand the potential mechanisms of recruitment of peripheral immune cells to the brain after LPS exposure, we used CellChat [49] to characterize signaling pathways between brain-resident microglia and peripheral immune cell subtypes. Considering first WT animals treated with LPS, monocytes and macrophages displayed the strongest outgoing and incoming signals, although all cell types had evidence of responding pathways [Fig 3A, B]. Some of the most strongly regulated pathways across cell types included those related to secreted signaling (CCL), proinflammatory PPIA signaling (CypA), and ECM-receptor interactions (THBS). We further examined which ligand-receptor pairs were signaling from microglia to other cell types and vice versa. Under LPS exposure, microglia signaled via CC chemokine ligands, which were received by multiple peripheral immune cell types via chemokine receptors [Fig. 3C]. Microglia also received diverse signaling from other cell types, including *ApoE* signaling to *Trem2* from macrophages and monocytes, complement C3 to C3 receptor (i.e., *C3* – *C3ar1*), and semaphorin to plexin (i.e., *Sema3a/d* – *Plxnb2*) signaling [Fig. 3D]. These results indicate that the model used here of single intrastriatal injection of LPS induces profound changes in inflammatory signaling, including cell to cell communication between brain-resident and recruited peripheral immune cells.

**Figure 3.**
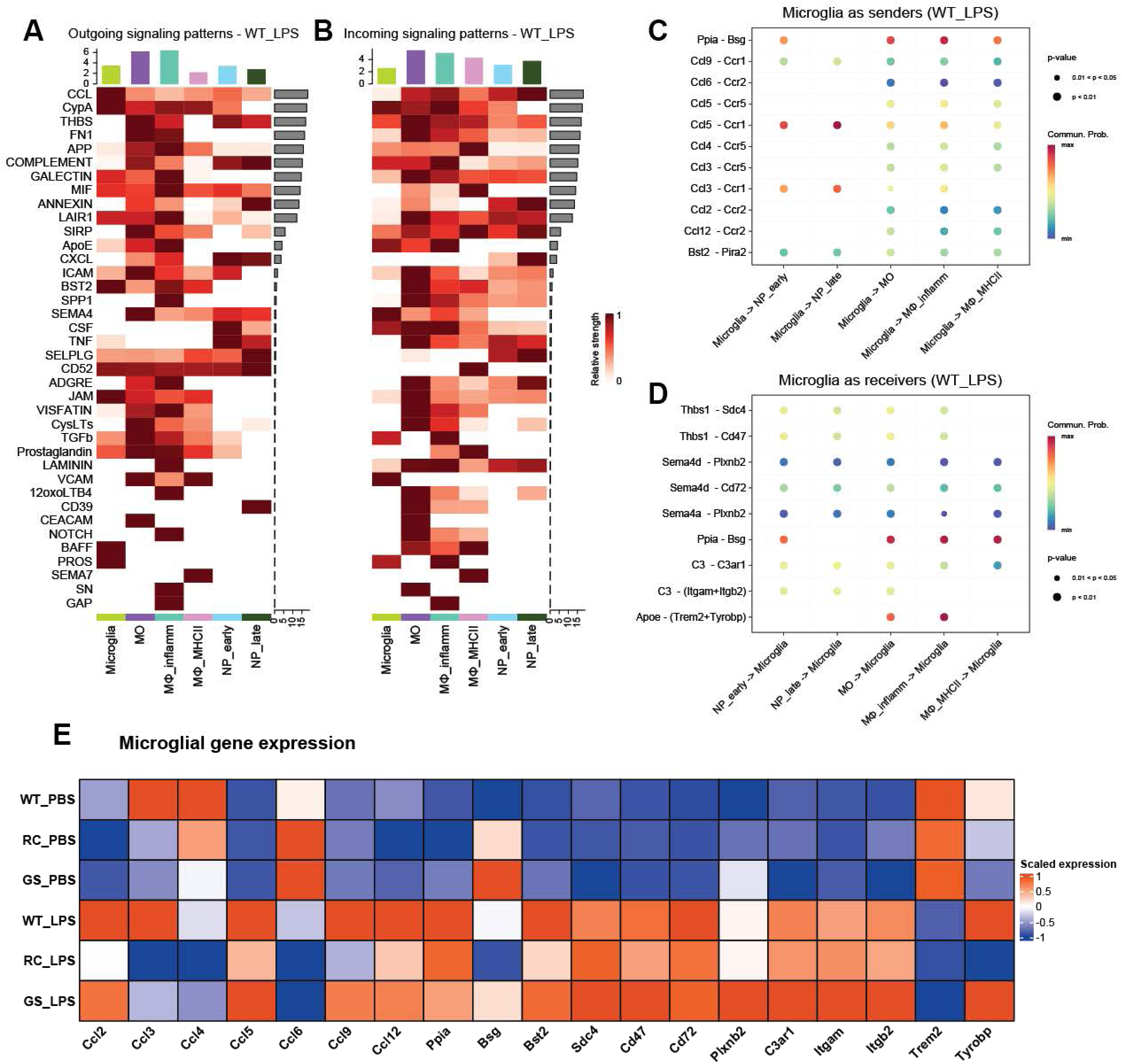
**Cell:cell communication genes associated with LPS injection.** [A, B]. Heatmaps of outgoing [A] or incoming [B] signaling patterns are colored by signaling strength for a given pathway (shown vertically) and cell type (shown horizontally; MG, microglia; NP, neutrophil; MO, monocytes; Mɸ, macrophage). Colored bars above the plot show cumulative signaling strength for a given cell type and gray bars to the right indicate summary of signaling strength for a given pathway across all cell types. [C, D] Bubble plots for ligand-receptor pairs with microglia as senders [C] or receivers [D] in LPS treated wild type mice. Ligand-receptor pairs with significant (p<0.01) interactions are shown vertically and cell pairs horizontally. Dots are colored by communication probability. [E]. Heatmap of scaled gene expression, as indicated on the color key on the right, for key genes identified in [C,D] along the horizontal axis for genotype - treatment pairs shown vertically.

### Limited effect of Lrrk2 genotype on LPS-induced gene expression

The above data establish that a single striatal injection of LPS resulted in profound changes in gene expression in both brain resident and peripheral immune cells, as expected. However, a primary objective in this study was to understand how Lrrk2 genotype, analogous to those causing human PD, influence gene expression patterns. We initially examined microglial gene expression in genes associated with cell-cell communication patterns [Fig 3A-D]. We found general upregulation of genes in the Ccl family and concomitant down regulation of *Trem2* in wild type animals after LPS exposure [Fig 3E]. However, Lrrk2 genotype showed no effect on expression of these genes under LPS stimulation [Fig 3E].

Given that these results were in opposition to our initial hypothesis that Lrrk2 mutations would exacerbate neuroinflammation, we looked more broadly at the gene expression patterns in the three different genotypes of mice in the immune populations examined previously. As expected, we observed that LPS induces substantial changes in gene expression in wild type microglia, with over 5000 differentially expressed genes (DEGs) between wild type and LPS treatment [Fig 4A; Table S7-10]. Some of the most substantially affected genes included known LPS-responsive genes, including *Saa3*, *Lcn2*, and *Arg1*, showing over 1000-fold increases. By contrast, we found minimal effect of Lrrk2 genotype on microglial gene expression in either PBS- or LPS-treated conditions, with only 2 genes (*Serpine2* and *Ccr5*) being significantly differentially expressed with a FDR < 0.1 in an interaction model between the R1441C genotype and LPS [Fig 4A].

**Figure 4.**
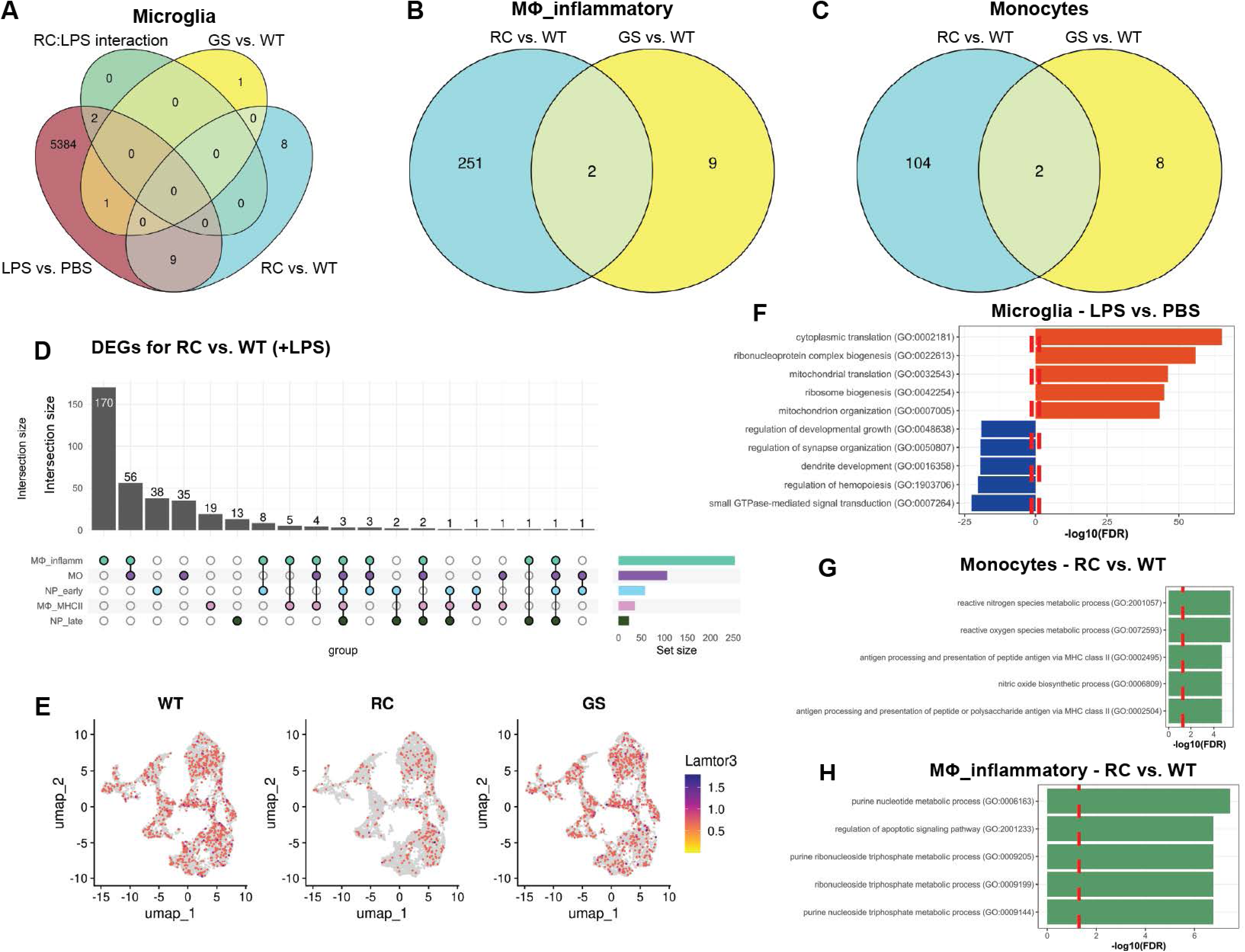
**Differential gene expression with LPS treatment and genotype** [A] Venn diagram showing overlapping differentially expressed genes (DEGs) between the effect of LPS, LRRK2 genotype, and their interaction in pseudobulked microglia (analysis in DESeq2 with formula: *counts ∼ genotype + LPS_treatment + genotype:LPS_treatment*; See Table S7-10). [B-C] Venn diagram showing overlapping DEGs between the effect of R1441C and G2019S in pseudobulked [B] macrophages (Tables S11-12) and [C] monocytes (Table S 13-14; formulae: *counts ∼ genotype*). [D] Upset plot for the number of DEGs in LPS treated R1441C knockin compared to LPS treated wild type animals in cells indicated vertically (MG, microglia; NP, neutrophil; MO, monocytes; Mɸ, macrophage; Table S15-19). Vertical bars show number of DEGs in indicated intersections while total numbers of DEGs per cell type are shown in hard on the right of the plot. [E]. Universal manifold approximate projections of expression of *Lamtor3* in LPS treated animals, from left to right wild type, R1441C and G2019S knockin. [F-H] Gene enrichments in immune cell types after LPS treatment. For gene sets along the vertical axes, −log10 P after FDR correction is shown for microglia in wild type animals comparing LPS with PBS [F; Table S20] and monocytes [G; Table S21] and macrophages [H; Table S22] comparing LPS treated R1441C knockin with wild type LPS treated.

We next examined differential expression comparing LPS-treated wild type and Lrrk2-mutant animals across all peripheral myeloid subtypes. Note that for these comparisons, we were unable to use a genotype:treatment interaction model due to the complete absence of these cells in the brain under PBS-treated conditions, so we opted to only test the effect of genotype within LPS-treated animals. Importantly, this approach does not allow for testing whether genotype differences are LPS-dependent or not. Across cell types, the R1441C genotype displayed more significant DEGs than G2019S relative to wild type, possibly due to a lower n for the G2019S + LPS condition [Fig. 4B, C; Table S11-14]. Given this, we examined differential expression in R1441C mutants further. We found the greatest number of DEGs in inflammatory macrophages and monocytes (253 and 106 DEGs, respectively), with substantial sharing between them [Fig. 4D]. 3 genes, *Amdhd2*, *Cap1*, and *Lamtor3* were differentially expressed across all cell types, [Fig. 4D; Table S15-19], suggesting that this is a consistent response to the R1441C mutation across immune cell types. However, there was not evidence that the same effect is present in response to the G2019S mutation [Fig. 4E].

Additionally, we further analyzed DEGs for functional enrichment in these different comparisons. Genes upregulated in microglia in response to LPS showed strong enrichment in translation, and downregulated genes were enriched for terms such as synapse organization and small GTPase-mediated signal transduction [Fig. 4F; Table S20]. Given the limited numbers of DEGs in R1441C vs. wild type comparisons for other immune cell subtypes, we opted to test for enrichments using all DEGs, regardless of up- or downregulation, to evaluate broad pathway-level dysfunction [Fig. 4G, H]. In monocytes, DEGs were enriched for terms including reactive oxygen and nitrogen species metabolism and MHC-II-related antigen processing and presentation [Fig. 4G; Table S21]. In inflammatory macrophages, DEGs were enriched for terms such as purine metabolism and regulation of apoptosis [Fig. 4H; Table S22]. Collectively, these results demonstrate that, in the context of a substantive transcriptional response of immune cells to LPS, Lrrk2 genotype has a much more modest effect.

## Discussion

A large body of literature now indicates that genetic risk factors play a substantive role in determining lifetime risk of PD [52,53]. In addition, it is established that aging also contributes to PD [54] along with non-genetic factors such as exposure to viruses [55]. One way in which these different risk factors may contribute to disease is via inflammation and, in this context, the LRRK2 gene is particularly interesting because it is expressed at high levels in macrophages and related cells, including brain-derived microglia [19,20,35,36]. We therefore aimed to examine how mutations in LRRK2 affect responses to neuroinflammation *in vivo* at single cell resolution. Surprisingly, we find that in a model with strong transcriptional responses to acute neuroinflammatory signaling across many cell types, the effects of endogenous Lrrk2 mutations are modest.

We were able to establish that the use of a single injection of LPS in the striatum evoked strong inflammatory responses across all cell types in the striatum and ventral midbrain. For example, we found multiple genes in microglia that responded to LPS injection including upregulation of the scavenger receptor *Msr1* [56] and downregulation of *Bin1*, a multifunctional protein proposed to regulate neuroinflammatory responses in microglia [57], both of which were validated using HiPLex RNAscope assays. Similarly, we found strong evidence of transcriptional remodeling after LPS exposure in other immune cells. Based on the relatively high number of lymphocytes and macrophages in the LPS compared to PBS treated animals, we infer that there is movement of peripheral immune cells towards the brain in this model, despite being restricted to a single CNS injection. Additionally, analysis of single cell gene expression data suggests interactions between different cell types for canonical immune signaling pathways. Therefore, the current model has expected and valid responses across multiple immune cells.

We additionally found evidence of inflammatory responses in other resident brain cells. For example, genes related to inflammatory response (*Ackr1*) or immune response through the complement system (*Cd93*) were upregulated upon LPS treatment. Conversely, downregulated genes were reported to be involved in membrane maintenance (*Utrn1*) or in lipid metabolism in ER and Golgi (*Sgms1*) and may influence blood-brain barrier permeability and vascular transport, all of which were validated. We noted an unusual decrease in astrocyte cell numbers in the single cell data. As staining of GFAP demonstrated an upregulation of astrocytes in LPS treated animals, similar to prior reports in literature [58] we infer that the cell recovery method used here likely underestimates activated astrocytes. We did not find evidence of neuronal cell death at the specific time point we chose after LPS injection, suggesting that any expression changes in immune cells are not related to consequences of neuronal loss. In general, expression changes in neurons were modest compared to other cell types discussed above, although we did confirm our previous findings [48] that upregulation of the Iron metabolism gene *Fth1* occurs in all cell types examined.

Collectively, these data show that monitoring transcriptional profiles at a single cell level can be used to confirm that there are robust responses to inflammatory signals *in vivo* using this model. However, the effect of Lrrk2 genotype were much more modest than the signal seen with LPS treatment, across all cell types identified. The lack of effect of Lrrk2 variants is in contrast to prior data showing, in BAC transgenic mouse models, a strong effect of genotype on responses to peripheral LPS injection [37,38]. There are multiple differences in experimental design between the present study and prior reports that may contribute to the alternate outcomes regarding Lrrk2 genotype. First, we used central rather than peripheral LPS exposure, which may limit our ability to detect responses that begin outside of the brain, although our observation that there is evidence of peripheral immune cell infiltration argues against this possibility. Second, we used a relatively acute exposure paradigm, which might be important if LRRK2 plays more substantive roles in recovery from inflammation rather than acute initial responses. Third, our study used endogenous mouse Lrrk2 variants, compared to overexpression of human LRRK2 in prior studies. At the current time it is not known if LRRK2 shows differential regulation in different species. Finally, we note that Kozina et al., used large scale measures of the proteome rather than transcriptome as required in our design to look at single cell level responses. Given that LRRK2 has known roles in protein trafficking and lysosomal function [25], one reasonable interpretation is that we would have needed to extend the current study to protein-level responses. In this context, it is potentially of interest that the sparse differential gene expression patterns between mutant and wild type animals include changes in genes related to endolysosomal function, which may represent compensatory mechanisms.

With the acknowledgement that any combination of the above factors may be at play, we propose that the current results should motivate additional investigations into the role of endogenous LRRK2 mutations in responses to neuroinflammation *in vivo*.

## Acknowledgements

We thank the NISC Sequencing Core Facility (National Institutes of Health (NIH),Bethesda) for their assistance with sequencing and technical support.

## Data Availability Statement

Single cell RNASeq data has been deposited to GEO (GSE265833). Scripts to perform the analysis as described in this manuscript are available on github (https://github.com/neurogenetics/Lrrk2_LPS_scRNAseq).

## Supplementary figure legends

**Fig. S1.**
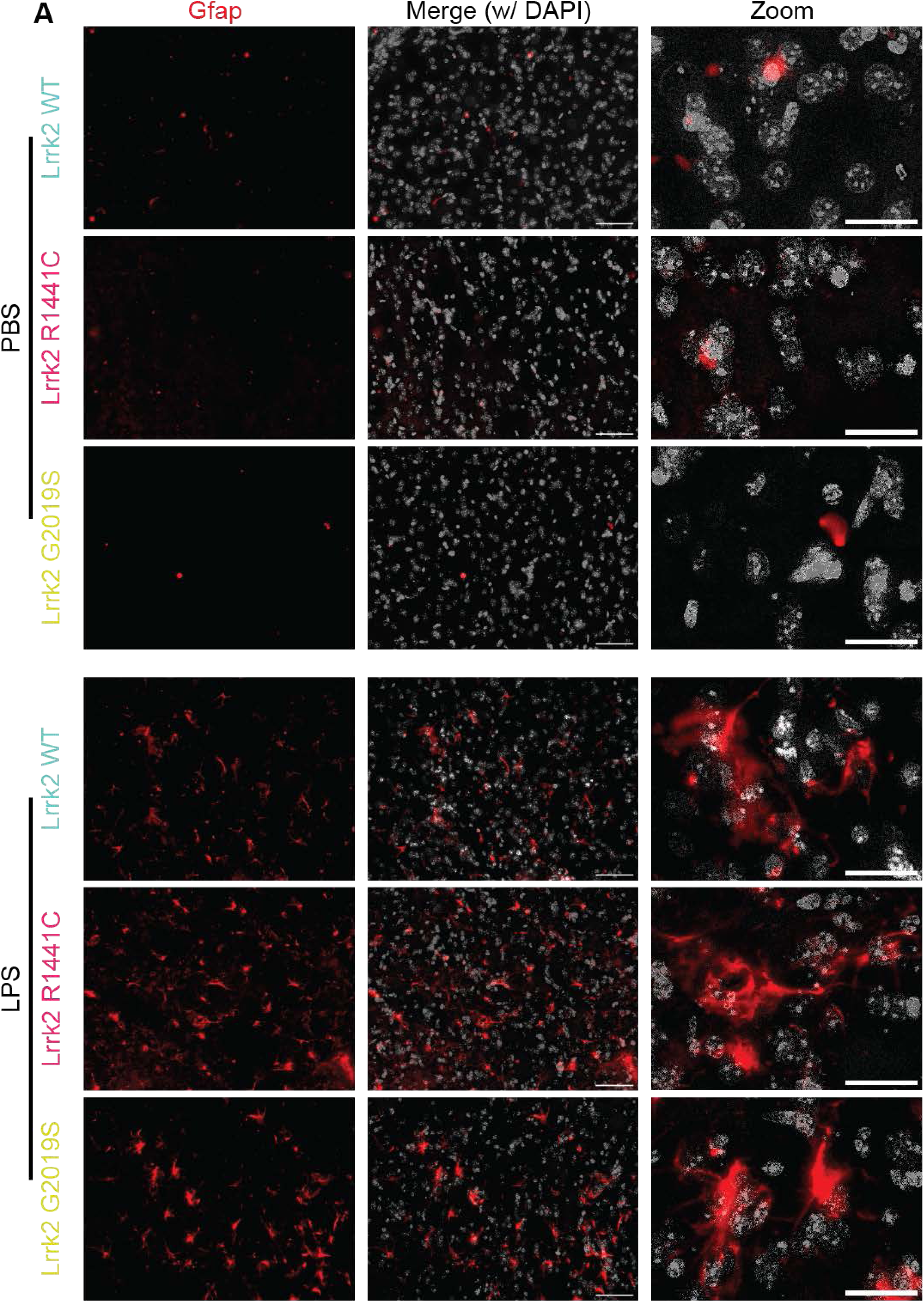
**LPS injection induces Gfap in striatal astrocytes *in vivo*.** Immunohistochemistry and confocal microscopy images of the striatum of PBS (upper panels) or LPS (lower panels) injected mice stained with rabbit Gfap-specific antibody and visualized with anti-rabbit Alexa 568 (red). Cell nuclei are stained with DAPI (gray). Representative examples are shown for wild type (WT), R1441C or G2019S knockin animals as indicated. Scale bar corresponds to either 50 μm (overview images) or 20 μm (zoomed images).

**Fig. S2.**
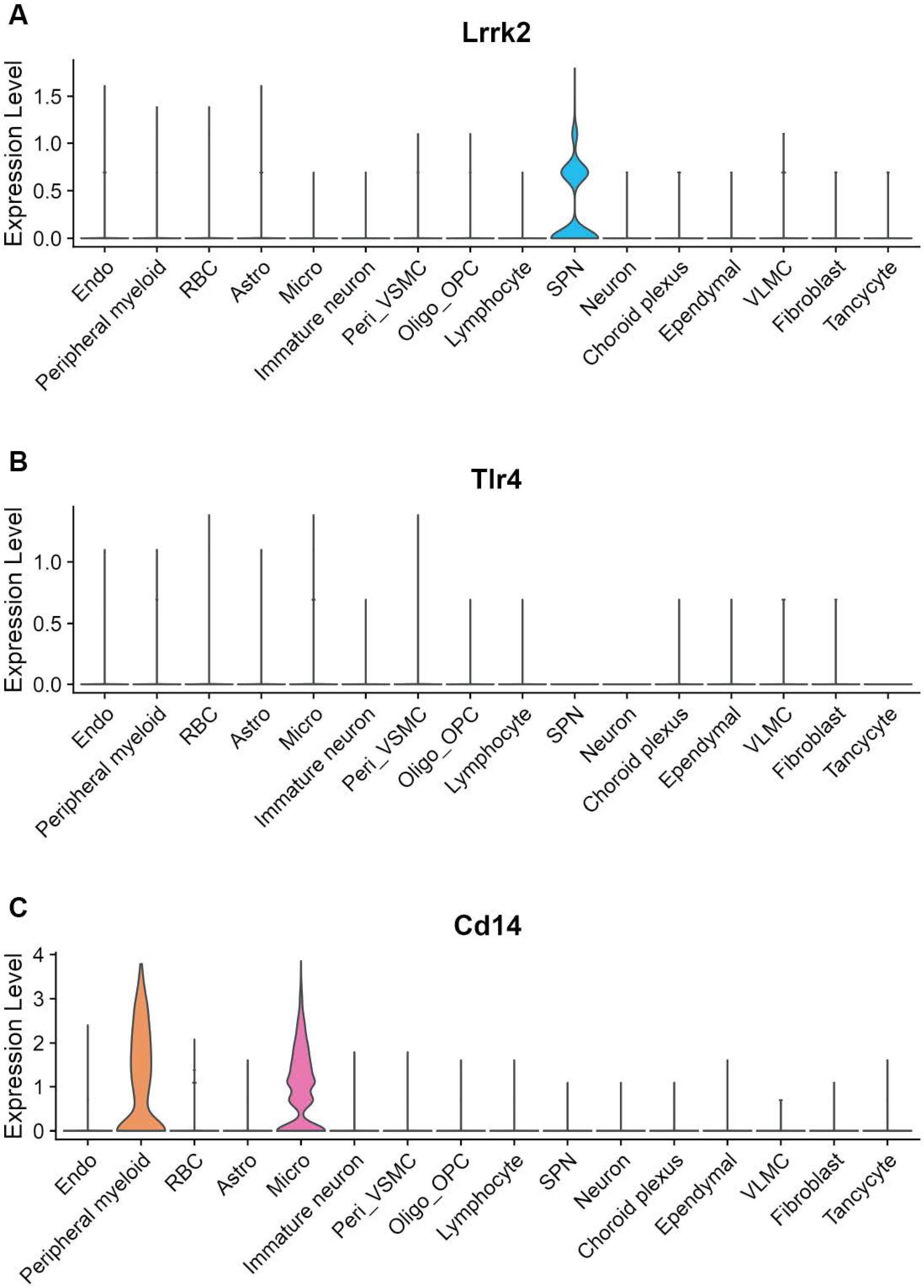
Gene expression of Lrrk2 and primary LPS receptors, Tlr4 and Cd14. Feature plots demonstrating expression patterns of *Lrrk2* (A), and of two primary LPS receptors, *Tlr4* (B) and *Cd14* (C) on microglia and macrophages. Universal manifold approximate projection (UMAP) positions are as in main figure 1. Relative expression levels are indicated by color intensity.

**Fig. S3.**
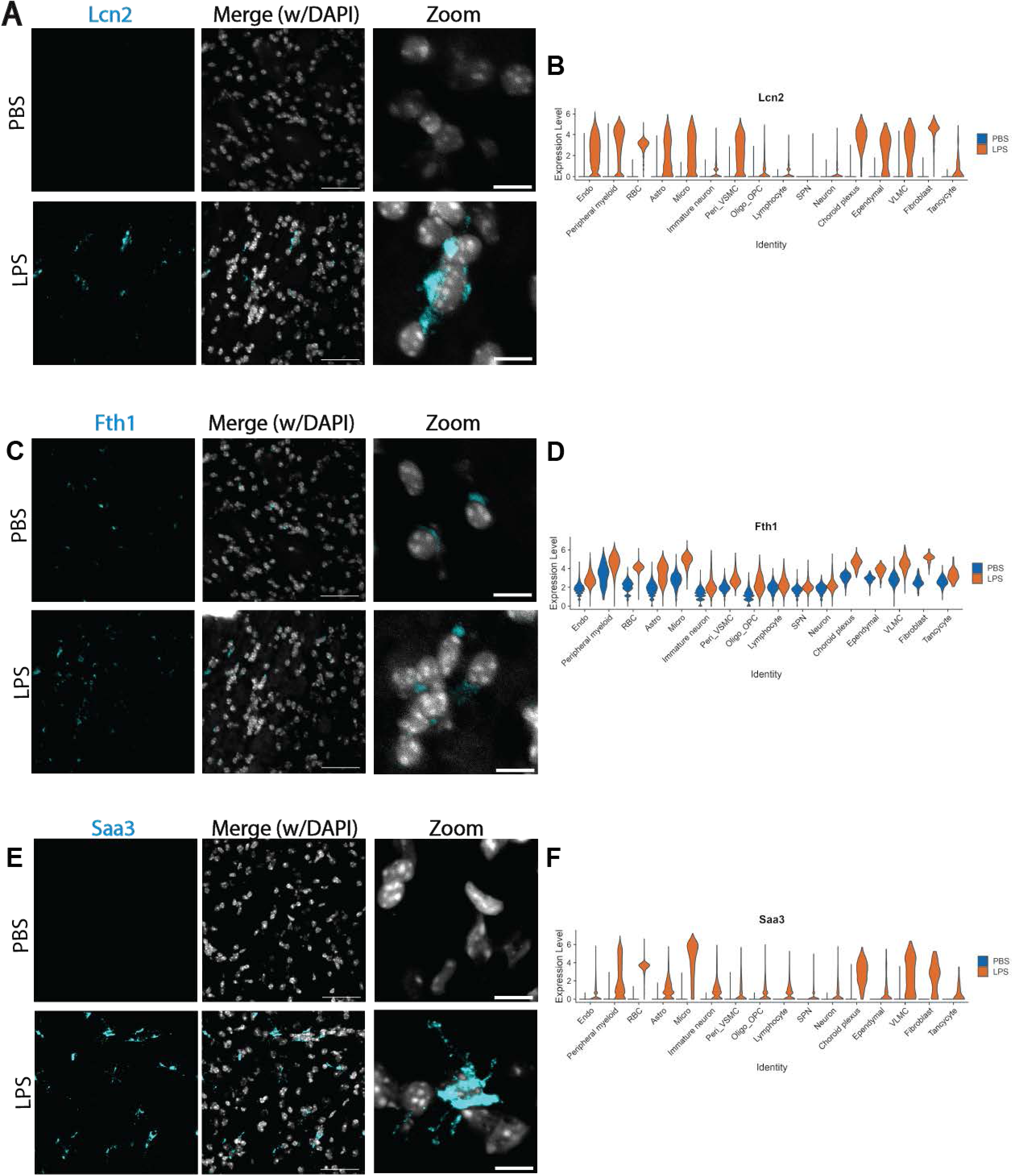
Validation of non-cell type-specific LPS-responsive expressed genes using RNA in situ hybridization. [A,C,E] Representative HiPlex RNAscope of *Lcn2* [A] and *Fth1* [C] from the same field in PBS (upper panels) or LPS (lower panels) WT animals. Similarly, representative HiPlex RNAscope of Saa3[E], in PSB and LPS. Scale bars are either 50μm (overview images) or 10μm (zoom image on the right). [B, D, F] Violin plots showing upregulation of relative expression of *Lcn2* [B], *Fth1* [D] or Saa3 [F] against cell types indicated along the horizontal axis genes in PBS (blue) or LPS (red) treated wild type animals.

**Fig. S4.**
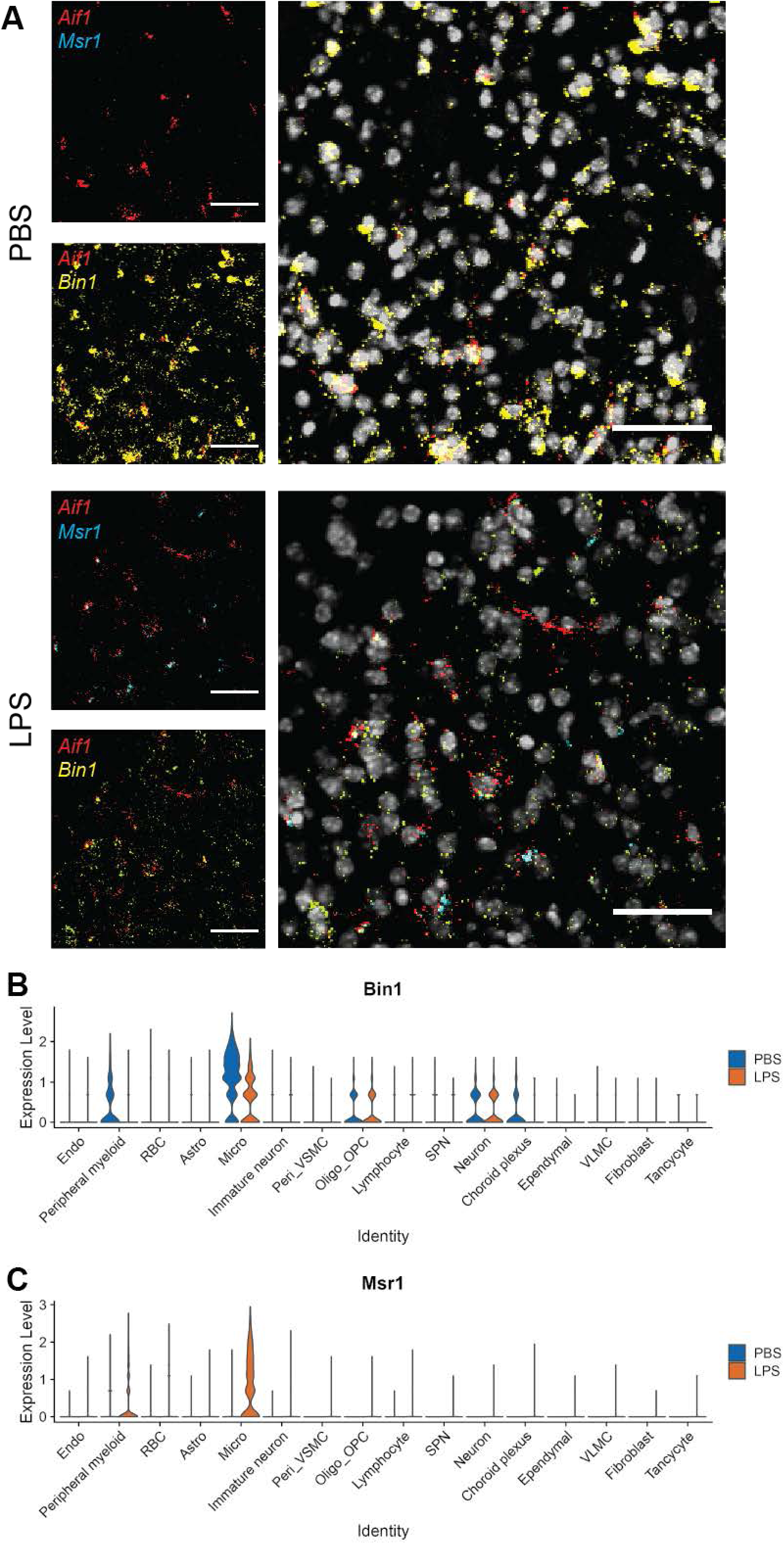
**Validation of cell type-specific LPS-responsive expressed genes using RNA in situ hybridization.** [A] Representative HiPlex RNAscope of simultaneous detection of *Aif1* (red), *Msr1* (blue) and *Bin1* (yellow), in PBS (upper panels) or LPS (lower panels) WT animals. Merge of all three channels is shown on the right. Scale bars represent 50μm. [B, C] Violin plots showing upregulation of relative expression of *Bin1* [B], or *Msr1* [C] against cell types indicated along the horizontal axis genes in PBS (blue) or LPS (red) treated wild type animals.

